# Modifying root/shoot ratios improves root water influxes in wheat under drought stress

**DOI:** 10.1101/2021.08.04.455065

**Authors:** Harel Bacher, Yoav Sharaby, Harkamal Walia, Zvi Peleg

**Author notes:** To whom correspondence should be addressed: Zvi Peleg, The Robert H. Smith Institute of Plant Sciences and Genetics in Agriculture, The Hebrew University of Jerusalem, Rehovot 7610001, Israel., Harkamal Walia, Department of Agronomy and Horticulture, The University of Nebraska-Lincoln, Lincoln, NE, USA.

## Abstract

Drought intensity as experienced by plants depends upon soil moisture status and atmospheric variables such as temperature, radiation, and air vapour pressure deficit (VPD). Although the role of shoot architecture with these edaphic and atmospheric factors is well-characterized, the extent to which shoot and root dynamic interactions as a continuum are controlled by genotypic variation is less known. Here, we targeted these interactions using a wild emmer introgression line (IL20) with a distinct drought-induced shift in the shoot-to-root ratio and its drought-sensitive recurrent parent Svevo. Using a gravimetric platform, we show that IL20 maintained higher root water influx and gas exchange under drought stress, which supported a greater growth. Interestingly, the advantage of IL20 in root water influx and transpiration was expressed earlier during the daily diurnal cycle under lower VPD and therefore supported higher transpiration efficiency. Application of structural equation model indicates that under drought, VPD and radiation are antagonistic to transpiration rate, whereas the root water influx operates as feedback for the higher atmospheric responsiveness of leaves. Collectively, our results suggest that a drought-induced shift in root-to-shoot ratio can improve plant water uptake potential in a short preferable time window determined by both water and atmospheric parameters.

## Introduction

Inadequate water availability is the major environmental factor limiting wheat productivity in many parts of the world. Drought episodes will become more persistent and more extensive, with projected climate change, and have numerous implications for rain-fed agricultural productivity (Gornall *et al*., 2010). Such drought stress is a consequence of not only precipitation distribution but also soil properties (*i.e*., texture and structure) and atmospheric variables (*i.e*., temperatures, relative humidity, and air vapour pressure deficit [VPD]) (Anderson *et al*., 1992; Rashid *et al*., 2018; Ihsan *et al*., 2016). Under such conditions, the plant’s ability to cope with the associated hydraulic constraints and maintain the soil–plant–atmosphere continuum (*i.e*., SPAC; Passioura, 1982) is mediated by various adaptive mechanisms. In general, water flows down potential gradients from the soil matrix to the leaf and evaporates through the stomata. Water stress can result in a breakdown of this continuum and may lead to hydraulic failure. Plants can regulate water loss by modulating the stomatal aperture as part of a short-term response to water limitation and through alteration of root system architecture as part of longer duration water stress (Sperry *et al*., 2002; Koevoets *et al*., 2016; Mansfield *et al*.,1990).

The root system architecture (*i.e*., root length and spatial distribution), as well as its anatomical and hydraulic properties, regulates plant water flow and maintains the whole plant water balance (Vandeleur *et al*., 2014; Sivasakthi *et al*., 2017). Root hydraulic conductance amplitudes follow the daily circadian rhythm which leads to higher conductance during the night and early morning when plant demand is lowest. During the afternoon, when plants lose their turgor and the evaporative demand is high due to atmospheric effects, plant conductance decreases significantly (Caldeira *et al*., 2014). The intensity of this diel pattern increases under water stress or high evaporative demand, which indicates the adaptive nature of this mechanism, optimizing water use efficiency.

The complex array of aboveground and belowground tissue interactions is strongly related to atmospheric diurnal variations. One of the main driving forces of transpiration rate is the VPD (Monteith, 1995; Lobell *et al*., 2014). In response to water stress, high VPD, or their combination, the plant will increase or limit its water use. While the first will continue to exhaust its water residuals, the second will reduce its stomatal conductance thereby decreasing transpiration and carbon fixation rate, and eventually shrinking the relative growth rate. These two strategies can be interpreted as phenotypic stability and plasticity responses, respectively (Nicotra *et al*., 2010; Bacher *et al*., 2021). Under drought stress, plants that exhibit plasticity responses by limiting their maximum rate of transpiration would show beneficial effects on productivity as a result of greater water-saving and higher transpiration efficiency (TE) (Tanner and Sinclair, 1983).

Under rain-fed agro-systems, wheat plants that can enhance their root system can exhaust the water residuals with higher efficiency (Golan *et al*., 2018; Voss-Fels *et al*., 2018). At the same time, smaller aboveground tissues will prevent excessive water loss due to lower shoot transpiration areas. Recently, we evaluated a large set of wild emmer wheat introgression lines (ILs) under contrasting water availabilities and identified a specific adaptive IL that harbours high phenotypic plasticity via a drought-induced shift in the root-to-shoot ratio mechanism (Bacher *et al*., 2021). Here, we tested this adaptive IL and its recurrent parent (Svevo), using a physiological approach of gravity lysimeters with water, soil, and atmospheric sensors. We aimed to dissect the interactions between these two genotypes water stress response mechanisms and their environmental cues. We hypothesize that the drought-induced alteration in the root-to-shoot ratio can support the water influx continuum under sub-optimal atmospheric conditions. Our specific aims were to ***i)*** characterize the physiological mechanism associated with whole-plant water flux, and ***ii)*** dissect the effect of genotypic-atmospheric–physiological interrelations on the water influx continuum under drought stress. Our results indicate that the IL’s adaptation to the atmospheric state under drought enhances transpiration efficiency. This is mediated by higher root influx during the early morning when VPD is low and is coupled to higher canopy conductance and growth.

## Material and methods

### Plant material

Two genotypes with contrasting drought responsiveness were selected for the current study, durum wheat cultivar Svevo (sensitive) and wild emmer wheat introgression line IL20 (adaptive). Svevo is an elite Italian durum wheat cultivar released in 1996 (CIMMYT line/Zenit) and considered as a reference for the quality and productivity of durum wheat. IL20 (BC3F5) consists of multiple (mapped) wild introgressions on five chromosomes (1A, 1B, 2A, 4A, 5B) accounting for ~4.5% of the accession Zavitan genome in Svevo. IL20 was selected from a set of ILs for its striking shift in the root-to-shoot ratio in response to water stress during the vegetative growth stage (Bacher *et al*., 2021).

### Lysimetric experiment-growth conditions

The experiment was conducted in the iCORE Center for Functional Phenotyping of Whole-Plant Responses to Environmental Stresses, the Hebrew University of Jerusalem, Rehovot, Israel, equipped with functional phenotyping gravimetric system, PlantArray 3.0 lysimeter system (Plant-DiTech, Rehovot, Israel). Before starting the lysimetric readings, uniform seeds of Svevo and IL20 were surface disinfected (1% sodium hypochlorite for 30 minutes) and placed in Petri dishes on moist germination paper (Anchor Paper Co., St. Paul, MN, USA) about 3 cm apart, at 24°C in the dark for 5 days. After that, uniform seedlings from each line were transplanted into cavity trays with potting soil, one seedling per cavity, and grown for seven days. Seedlings were then transplanted to 4-liter pots (one seedling per pot) filled with 2.8 kg fine silica sand 75-90 (Negev Industrial Minerals Ltd., Israel). Each pot was immersed in a plastic container through a hole in its top cover. Evaporation from the containers and pots was prevented by a plastic plate cover, punched in the middle. Pots were placed under semi-controlled temperature conditions, with natural day length and light intensity (Supplementary Fig. S1). Each pot was placed on a load cell with a digital output generated every three minutes (Dalal *et al*., 2020). Each plant was irrigated by four on-surface drippers to ensure uniform water distribution in the pots at the end of the irrigation event and before drainage. Plants were irrigated in three consecutive cycles, between 23:00 and 02:00. The daily predawn pot weight was determined as the average weight between 05:00 and 05:30 h after ensuring that all excessive water was drained from the pot (Dalal *et al*., 2020).

The temperature and relative humidity in the greenhouse and the photosynthetically active radiation were monitored using an HC2-S3-L Meteo probe (Rotronic, Crawley, UK) and LI-COR 190 Quantum Sensor (Lincoln, NE, USA) and a soil moisture sensor (5T, Decagon Devices, Pullman, WA, USA). Altogether, pots weight and environmental information are recorded by the sensors every 3 min and the data could be viewed in real-time through the online web-based software SPAC-analytics (Plant Ditech, Rehovot, Israel).

### Lysimetric experiment-experimental design

Plants were grown under optimal conditions for 25 days (12 to 32 in Zadoks growth scale; Zadoks *et al*., 1974) and then divided into two groups, well-watered (WW) and drought stress (DS) treatments, that were gradually applied over nine days. Under the DS, each plant was irrigated individually based on a 20% reduction from its previous day of transpiration, so that all plants were exposed to controlled drought treatment according to individual plant water demands. All plants were harvested after 35 days at the booting stage (45 in Zadoks growth scale) and oven-dried for 8 days at 80°C for dry weight measurements. Volumetric water content (VWC) of both genotypes under a specific water treatment was measured by using time-domain reflectometry (TDR) (5TE; Decagon Devices, Pullman, WA, USA).

### Whole-canopy weight and gas exchange measurements-lysimetric experiment

A full description of the physiological measurements methodology for the lysimetric system was described previously (Halperin *et al*., 2017). In brief, *whole-canopy daily transpiration* was calculated as the difference between the load-cell at pre-dawn (W_m_) and in the evening (W_e_) that were measured at 05:30 and 19:00, respectively:

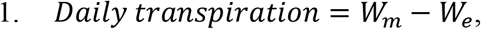

W_m_ and W_e_ values were determined as the average load-cell reading of the specified time to eliminate the effect of temporal variation in ambient conditions.

The *plant weight gain* was calculated post-irrigation at 05:30 after drainage had ceased. This enabled determination of plant weight gain (fresh weight biomass) for any desired period. The plant daily weight gain (Δ*PW_n_*) between consecutive days was calculated by:

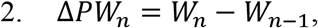

where W*_n_* and W*_n-1_* are the container weights after drainage on the following days, *n* and *n–1*, respectively.

The *whole-plant water-use efficiency* (WUE_w_) during the well-watered period was determined by the ratio between the sum of the daily plant fresh-weight gain (Δ*PW*) and water consumed throughout this period:

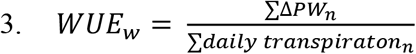

The WUE_w_ for the whole plant replaces the commonly used physiological WUE determined as the ratio between the accumulated CO_2_ molecules and evaporated H_2_O. Determination of plant weight on a single day (*PW_n_*) throughout the drought period was calculated differently from equations #2 and #3. Since the pots did not drain and plant weight gain could not be separated from the container and the watering scheme, the following calculation was made:

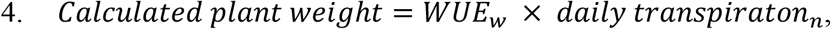

where *WUE_w_* was calculated for the well-watered period and *daily transpirationn* is the daily water consumption throughout the day (n). *Calculated plant weight gain* was determined by subtracting *calculated plant weight* from two consecutive days.

The *root water influx* was calculated by:

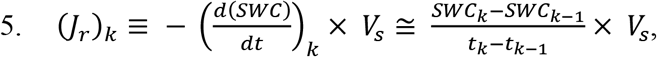

where W_k_ and W_k-1_ were the readings at time t_k_ and t_k-1_, respectively. Soil water content (SWC) in a pot was measured by the soil moisture sensor (5TE; Decagon Devices, Pullman, WA, USA), and *Vs* was the soil volume in the pot.

The *whole canopy conductance* (*gs_c_* [mmol sec^−1^ m-^2^]) was calculated by:

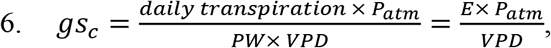

where *P_atm_* was the atmospheric pressure (101.3 kPa). VPD was determined as the difference (in kPa) between saturation vapour pressure and actual vapour pressure of ambient air. Transpiration rate was normalized by leaf weight (E [mmol sec^−1^ m-^2^]).

### Phytotron experimental design and conditions

Uniform seeds of Svevo and IL20 were placed on a moist germination paper (25 × 38 cm; Anchor Paper Co., St. Paul, MN, USA), about 2 cm apart, with the germ end facing down (*i.e*., Cigar Roll method; Watt *et al*., 2013), and placed in a growth chamber (16°C). Four uniform seedlings were transplanted into pre-weighed 4L pots and thinned to 2 plants per pot after establishment. The pots consisted of a soil mixture of 80% sandy loam and 20% peat. The pot surface was covered by a layer of vermiculite to prevent water evaporation from the pot surface. The experiment was conducted in the phytotron facility at the Faculty of Agriculture, Food and Environment, The Hebrew University, Israel. From seedling to tillering stage (23 in Zadoks growth scale) plants were grown under short-day (8/16h) at 16/12°C (day/night) and then transferred to 22/16°C (day/night) conditions under natural sunlight. A complete randomized experimental design consisted of two genotypes (Svevo and IL20) and two water treatments (WW and DS), with 6 replicates for each combination (total 24 pots). Up to the tillering stage, all plants were grown under well-watered conditions. Under WW treatment, pots were irrigated above actual consumption, to allow some water to drain. Under DS treatment, drought was initiated at tillering stage and pots were irrigated with 50% of the actual water consumption of the WW treatment. Fertilization was utilized once in ~10 days during the water application, using 0.1% soluble fertilizer [20/20/20 (NPK)] including micro and macro elements (Ecogen, Jerusalem, Israel). At the end of the experiment, plants were harvested and vegetative dry matter (DM) (stems and leaves) was separated from reproductive DM (spikes). Both tissues were oven-dried at 80°C and 38°C, respectively for 120 h and weighed. Spikes from each pot were then threshed manually to determine grain yield (GY).

### Gas exchange measurements

Plants were characterized for gas exchange parameters from the third leaf to grain filling (13, 21, 23, 33, 43, 65, and 85 in Zadoks growth scale) using a portable infra-red gas analyzer (LI-6800XT; Li-Cor Inc., Lincoln, NE, USA) to obtain the carbon assimilation rate (A), transpiration rate (E), stomatal conductance (gs), and total conductance to water vapour (g_tw_). Measurements were conducted at the mid-portion of the youngest fully expanded leaf or flag leaf in the later phenological stages (*n*=5). Photosynthetic photon flux density was set to 1500 μmol/m^2^ and the CO_2_ reference concentration was set as the ambient (400 ppm). The seasonally average leaf water use efficiency (WUE_*l*_) was calculated as the slope of the linear regression between A and E for each genotype within water treatment. A parallelism test was used to statistically differentiate between the curves representing genotypes under a specific water treatment.

### Measurements of leaf stomatal density

Stomatal density was determined using the rapid imprinting technique (Reich, 1984). Nail polish was applied to the adaxial side of the leaf, dried for 1 min, and then removed. Once it dried, the nail polish imprints were plated on glass coverslips and photographed under an inverted microscope (Zeiss, Jena, Germany). Imprints from each genotype were sampled (0.54 × 0.37 cm width and length) in four biological and three technical replicates. The technical replicates represented different leaf parts within one leaf sample that had been manually counted for stomata number.

### Statistical analyses and modelling

The JMP^®^ ver. 15 statistical package (SAS Institute, Cary, NC, USA) was used for statistical analyses unless specified otherwise. The longitudinal response was fitted for genotypes under each water treatment. Analysis of variance (ANOVA) was used within each day of the experiment to assess the first day of significant differences between the genotype within water treatment at the relative growth rate and daily transpiration dependent variables. Hourly basis graphs of normalized transpiration rate (E), whole canopy conductance (gs_c_), and root influx have been analysed within an hour on a specific day and tested for differences between genotypes in a t-test. The maximum canopy conductance and maximum root influx were determined from the mean of the last five days of maximum values within an hour (*n*=5). The differences between genotypes were tested on an hourly basis with a t-test within water treatment.

The differences in gas exchange parameters: Assimilation rate (A), stomatal conductance (gs) and total conductance to water vapour (g_tw_) between the two genotypes under specific water treatment were tested using a t-test. The analysis was made from the booting stage (43 in Zadoks growth scale), similarly to the lysimetric final phenological stage (*n=5*). Genotypic differences in leaf water use efficiency (WUE_*l*_) under specific water treatment throughout the season were obtained by fitting a linear curve between A and E and applying a parallelism test between the two lines.

Structural equation model (SEM) analysis was performed using the JMP^®^ ver. 15 statistical package and the package “Lavan” (Oberski, 2014) under RStudio 3.5.1 (RStudio Team, 2020) to test the dependent and independent variables associated with Genotype-atmospheric-water availability. For testing the model accuracy, we used the Chi-square (χ^2^) test and Bentler’s comparative fit index (CFI), which provides additional guidance for determining model fit. Values greater than 0.90 are considered a good fit for the model (Bollen and Long, 1993; Hu and Bentler, 1999). The root mean square error of approximation (RMSEA), provides additional guidance for determining model fit, as values smaller than 0.10 are preferred (Bollen and Long, 1993; Hu and Bentler, 1999).

## Results

### The higher root-to-shoot ratio of IL20 supports a greater growth rate under drought stress

We have identified a wild emmer introgression line (IL20) with unique adaptive characteristics of drought-induced root-to-shoot ratio alteration under water stress during the vegetative period (Bacher *et al*., 2021). In this study, we used a high-throughput gravity lysimeter system to further characterize IL20 under drought stress (DS), initiated during the developmental transition from vegetative to reproductive stage. Twelve-day-old plants (12 in Zadoks growth scale; Zadoks *et al*., 1974) were grown under well-watered (WW) treatment for 25 days. Then, nine days of increasing drought stress treatment were applied, for each plant individually, to normalize their water stress as expressed in volumetric water content (VWC) (Fig. 1A).

**Figure 1.**
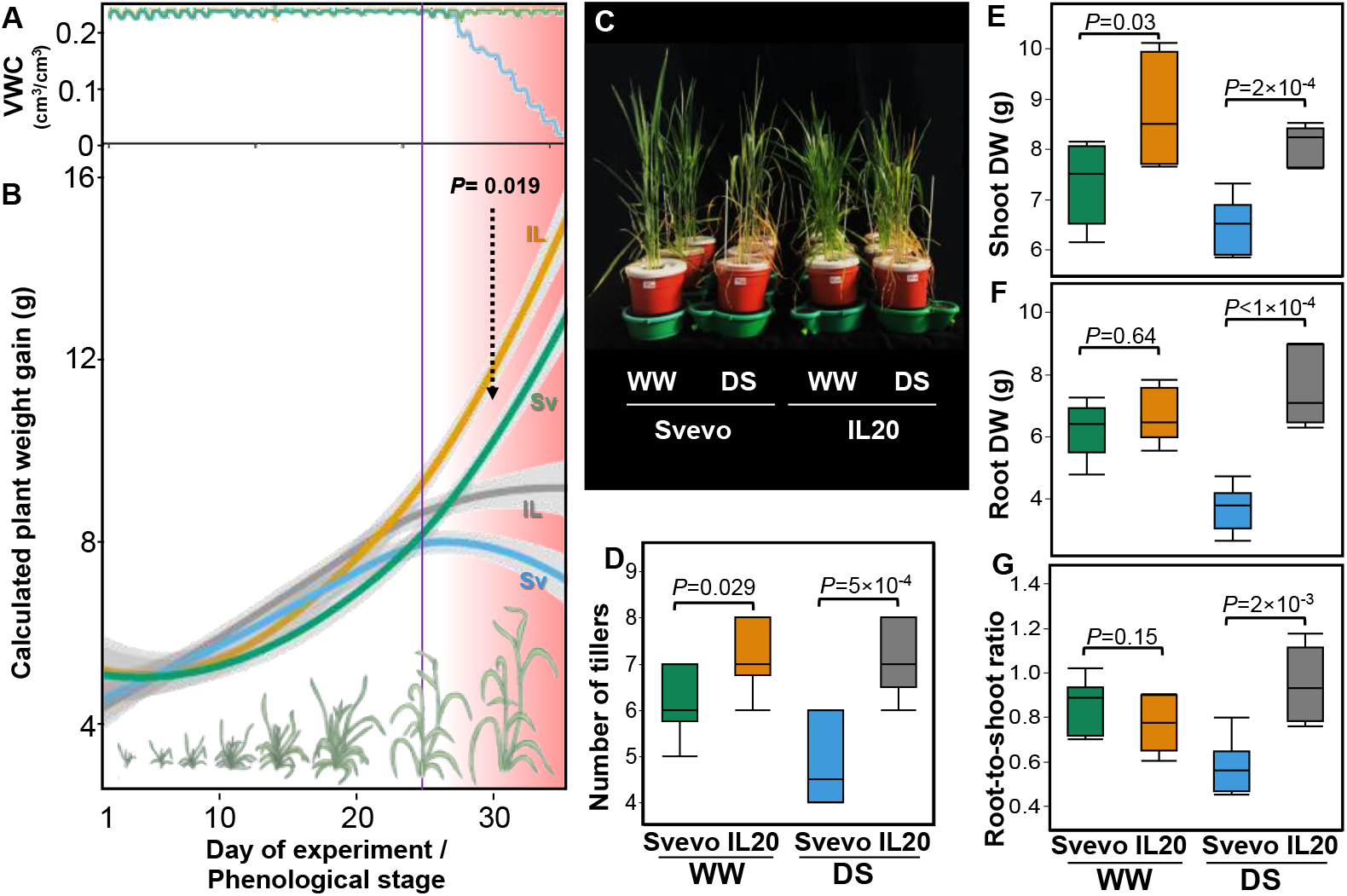
Longitudinal weight gain and its distribution under contrasting water treatments. (**A**) Volumetric water content (VWC) and (**B**) calculated weight gain dynamics of Svevo (Sv) and IL20 (IL) under well-watered (WW) and drought stress (DS) treatments along with the different wheat phenological stages. The vertical purple line represents the point that drought stress application started. The increasing stress intensity is represented by the pink background colour intensity. The line curves represent the genotype mean within water treatment and day (*n*=5). Shaded colours indicate the standard error. (**C**) Svevo and IL20 under contrasting water treatments on the last day of the experiment. (**D**) Number of tillers, (**E**) shoot dry weight (DW), (**F**) root DW, and (**G**) root-to-shoot ratio of the two genotypes under contrasting water treatments. Differences between genotypes within water treatment were analysed using a t-test (*n*=5) on the last day of the experiment (35 days after transplanting).

In general, under WW treatment IL20 exhibited higher calculated weight gain compared with Svevo, although it was not significant (*P*≤0.05) for every day (Supplementary Table S1). After initiating the water stress treatment (26^th^ day), IL20 maintained its growth rate over time as the stress intensity increased (Fig. 1B). The major divergence in calculated weight gain started at the stem elongation stage (34 in Zadoks growth scale; day 30) five days after the stress stated (*P*=0.019; Fig. 1B; Supplementary Table S1). IL20 maintained a lower and steady growth pattern (30-34 days) while Svevo presented a rapid decline in growth in response to water stress.

At the end of the experiment (booting stage; 45 in Zadoks growth scale), Svevo showed strong visual symptoms of leaf rolling and leaf senescence, while IL20 showed milder symptoms (Fig. 1C). The plants were harvested and shoot and root dry weight (DW) were analysed. IL20 exhibited a significantly higher number of tillers and shoot DW under WW treatment (*P*=0.029, and *P*=0.032, respectively) compared to Svevo, and under DS (*P*=0.0005, and *P*=0.0002, respectively) (Fig. 1D, E). Root DW of IL20 was similar under WW and two-fold higher under DS (*P*≤1×10^−4^) as compared with Svevo (Fig. 1F). Consequently, under DS IL20 exhibited a significantly higher root-to-shoot ratio (*P*=1×10^−3^; Fig. 1G; Supplementary Table S2).

### IL20 exhibited higher whole canopy transpiration under drought stress

Under WW conditions, biomass accumulation is associated with a higher gas exchange rate (*i.e*., assimilation rate, transpiration rate, and stomatal conductance). However, under drought stress conditions, a relatively higher gas exchange rate may not directly translate into increased biomass accumulation. Using the gravity lysimeter system we were able to trace the genotypic response to DS by measuring daily transpiration and growth curves. IL20 exhibited significantly higher daily canopy transpiration compared with Svevo, as differences in the transpiration between the genotypes increased together with the increment of water stress intensity (Fig. 2A; Supplementary Table S3). Notably, daily transpiration dynamics under DS showed a similar pattern as found for calculated weight gain (Figs. 1B and 2A). Since a larger canopy is associated with greater transpiration, we normalized the transpiration to the plant weight (E; g water/g plant). We tested the daily E patterns for each genotype between water treatments and observed that, under DS, IL20 was able to maintain higher E than Svevo as water stress intensified. Svevo showed a decline during the last five days of severe water stress (days 30-34). Notably, under WW treatment the E pattern of both genotypes was comparable in the experiment window (Fig. 2B, C).

**Figure 2.**
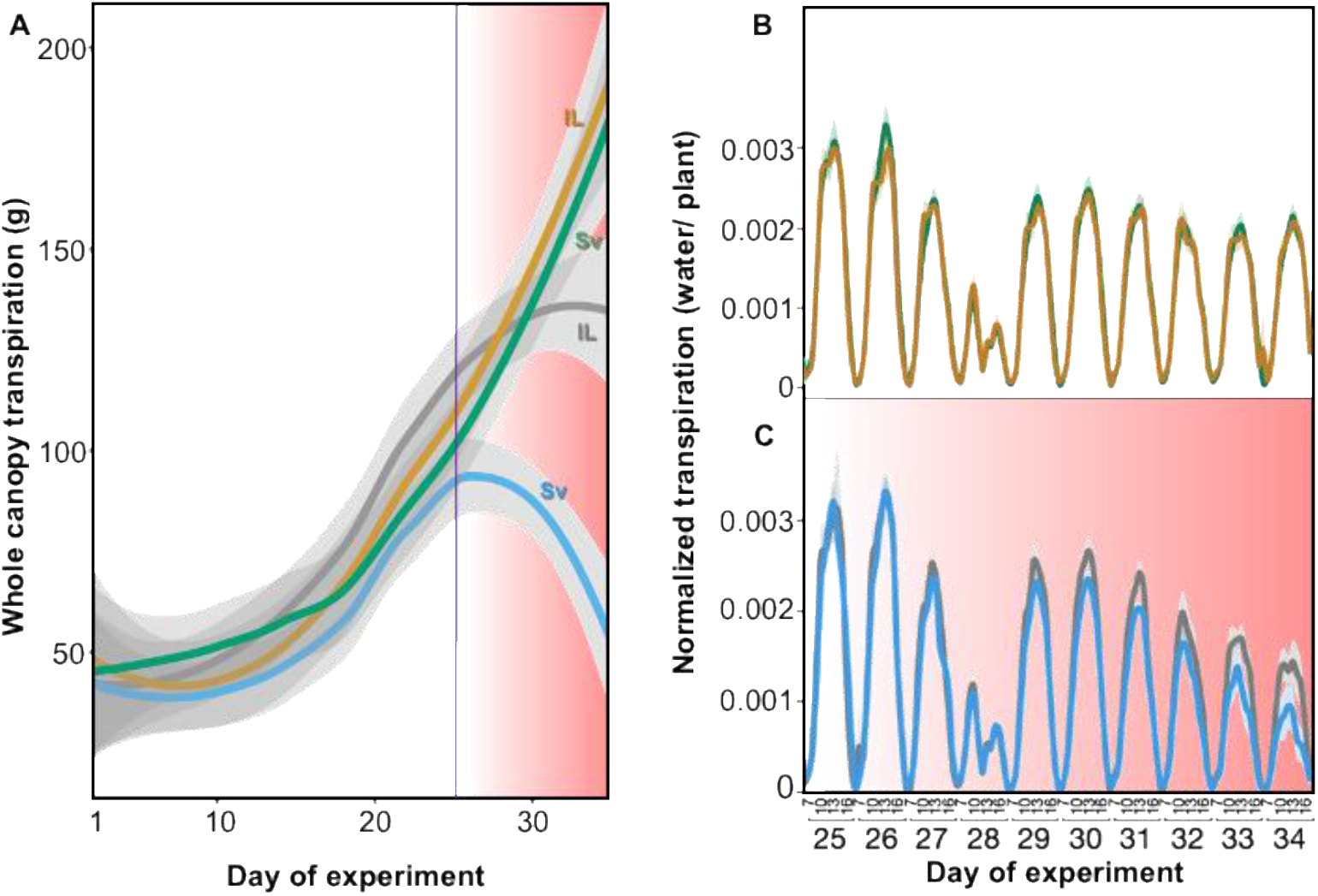
Whole canopy transpiration dynamics of Svevo (Sv) and IL20 (IL) under well-watered (WW) and drought stress (DS) treatments. (**A**) Daily transpiration dynamics along with the experiment. The increasing drought intensity is represented with the pink background colour intensity. The vertical purple line represents the point that drought stress application started. Hourly dynamics of normalized transpiration rate of Svevo and IL20 under (**B**) WW and (**C**) DS. The line curves represent the genotype means (*n*=5) within water treatment, and shaded colours indicate the standard error.

### IL20 exhibited a higher and longer period of whole canopy conductance

To better understand the factors contributing to higher E values under DS of IL20 compared with Svevo, we characterized flag leaf stomatal density under two water treatments. We hypothesized that higher stomatal density can be directly associated with higher E. In contrast, the results indicated that the genotypes had similar stomatal density under both water treatments and that the water stress did not have any effect (Supplementary Fig. S2). Given no stomatal density differences, we hypothesized that there might be a difference in stomatal conductance in response to atmospheric parameters (Radiation and VPD) that may increase the transpiration use efficiency of IL20. To test this hypothesis, we focused on whole canopy conductance (gs_c_) on an hourly scale during the day. Our results suggest that under DS, gsc of both genotypes peak before maximum radiation, with IL20 exhibiting significantly higher gsc compared with Svevo along with the increased stress intensity (Fig. 3A). Moreover, the gap between IL20 gs_c_ compared with Svevo gets bigger as water stress intensity increases. For example, on the 29^th^ day (4 days after initiation of DS), the genotypic differences in gs_c_ were found between 10:00-12:00 while on the 34^th^ day the differentiation started as early as 07:00 and lasted until 14:00. To further explore the hourly gs_c_ dynamics, we focused on the last five days of the experiment (days 29-34) when the water stress was the most severe and calculated the hourly maximum gs_c_ within each day. This analysis yielded genotypic differences in diurnal gs_c_ under DS. While under WW treatment there were no differences between the genotypes, under DS IL20 exhibited ~50% higher gs_c_ capacity compared with Svevo between 08:00-09:00 AM. In addition, most of the gs_c_ differences between the genotypes occurred in the morning time (07:00-10:00 AM), when VPD is low (Fig. 3B-C).

**Figure 3.**
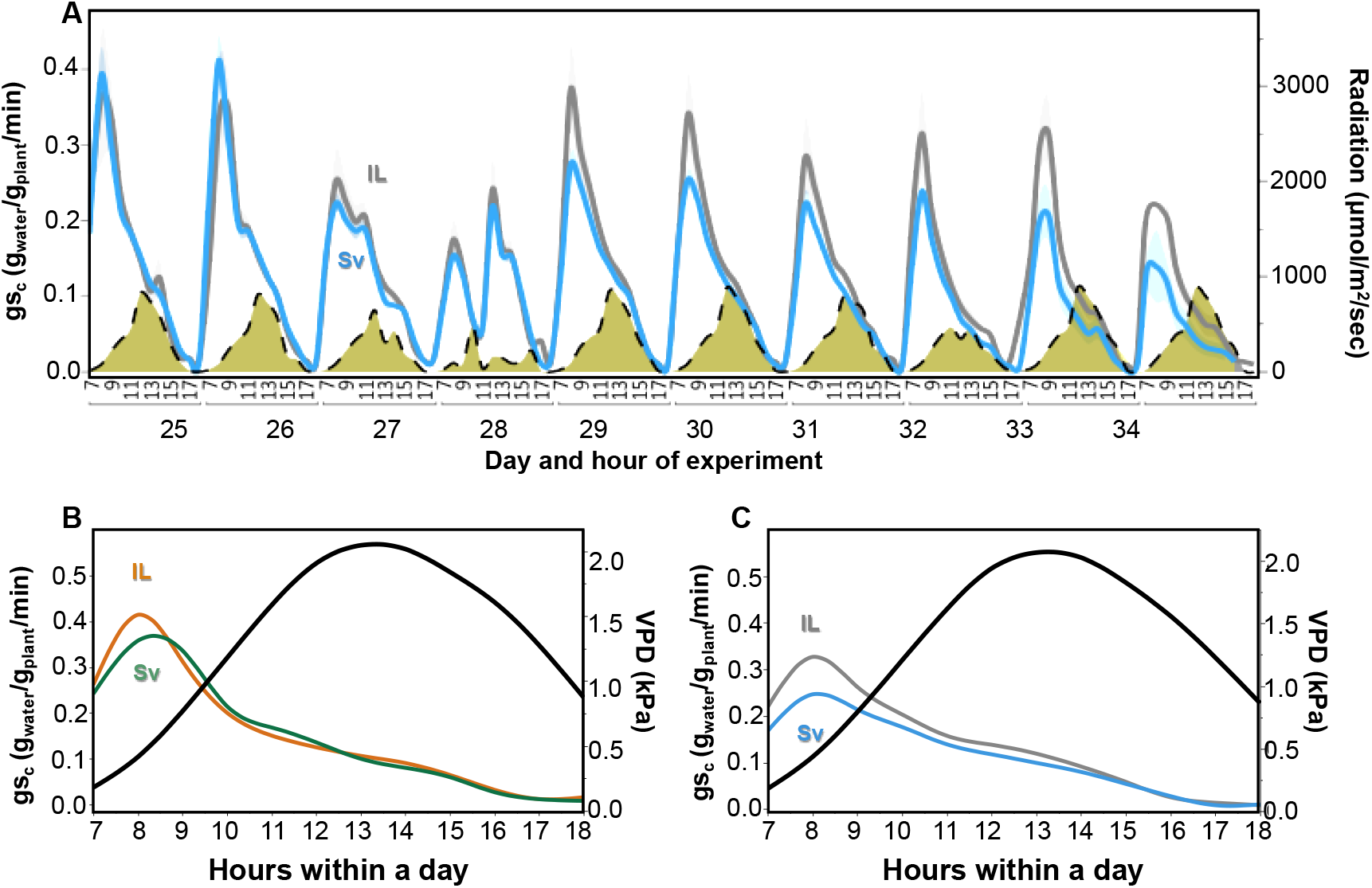
Diurnal dynamics of whole canopy conductance (gs_c_) of Svevo (Sv) and IL20 (IL) under well-watered (WW) and drought stress (DS) treatments. (**A**) Svevo and IL20 hourly gs_c_ dynamics together with the natural light radiation curve (yellow). The increasing drought intensity is represented with the pink background colour intensity. The maximum hourly gs_c_ average was calculated from the last five days (30-34) of the experiment under (**B**) WW and (**C**) DS treatments. The hourly vapour pressure deficit (VPD) is represented by the black curve. The genotype curves represent the genotype means (*n*=5) within water treatment, day, and hour. Shaded colours indicate the standard error.

### IL20 exhibited a higher gas exchange rate and leaf water-use efficiency under drought stress

To better understand the gs_c_ diel dynamics under DS, we tested the leaf gas exchange rate and extracted the leaf water use efficiency (A/E; WUE_*l*_) for both genotypes. We hypothesized that IL20 will exhibit higher WUE_*l*_ in response to water stress. Under both water treatments, IL20 gas exchange was higher compared with Svevo at the booting stage (this growth stage was the same stage we ended the lysimetric experiment). IL20’s higher gas exchange rate significance range was between *P*=0.03-0.06 (Fig. 4A, B, Supplementary Table S4). In addition, the seasonal average WUE_*l*_ of IL20 was higher than Svevo under DS (*P*≤1×10^−4^) (Fig. 4C, D, Supplementary Table S5) as total conductance to water vapour (gtw) was higher than Svevo under both water treatments (Supplementary Fig. S3). At the end of the season, IL20 vegetative dry matter (DM) and grain yield were higher than Svevo under DS (*P*=0.038, *P*=0.053, respectively) indicating that the advantage of higher gas exchange and WUE_*l*_ is associated with higher productivity (Fig. 4E, F).

**Figure 4.**
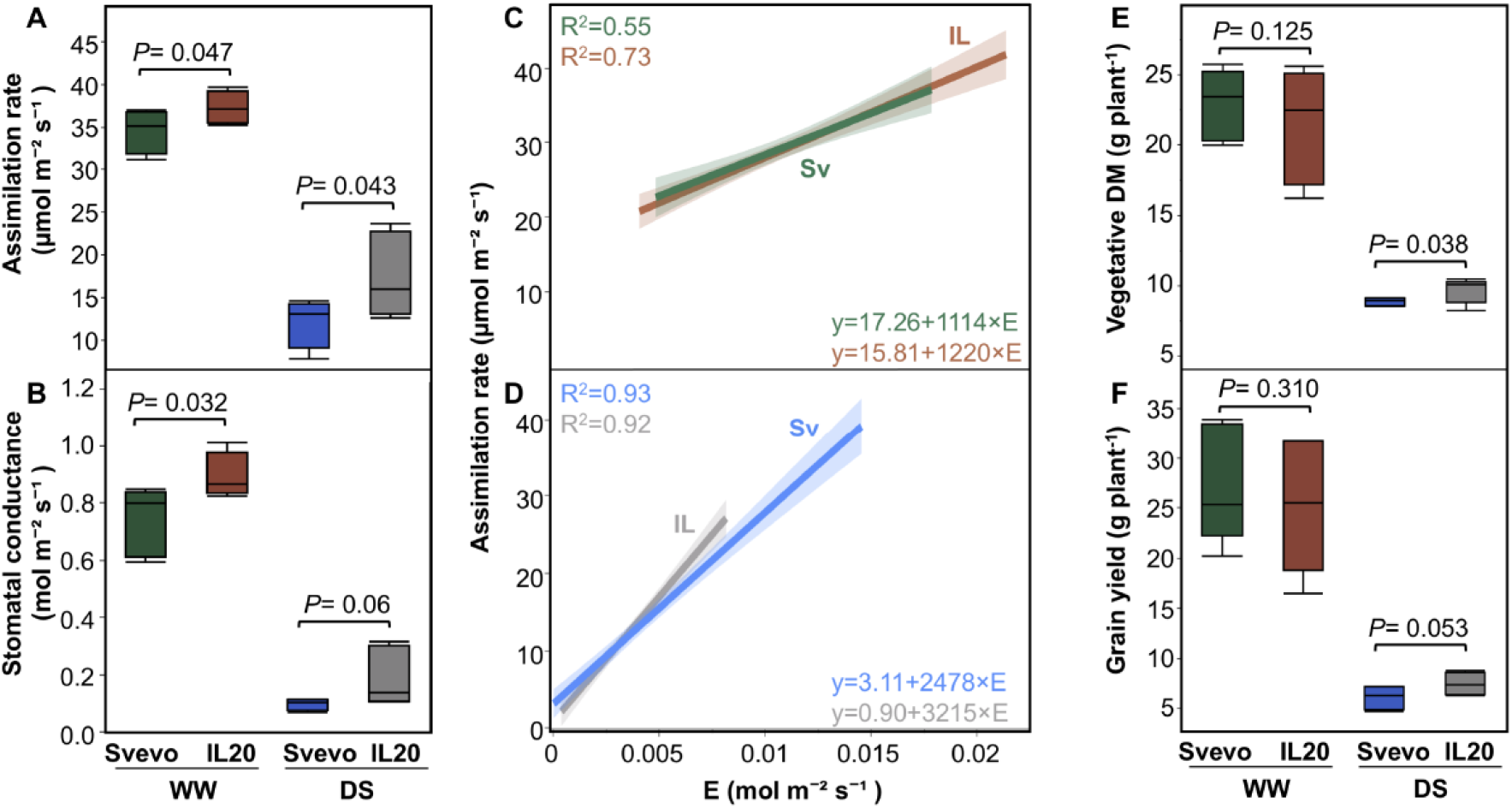
Leaf gas exchange, water-use efficiency and productivity of Svevo (Sv) and IL20 (IL) under well-watered (WW) and drought stress (DS) treatments. (**A**) Assimilation rate during the booting developmental stage. (**B**) Stomatal conductance during the booting developmental stage. Seasonal average of leaf water-use efficiency (A/E) under (**C**) WW and (**D**) DS treatments. (**E**) Vegetative dry matter (DM) and (**F**) grain yield. Differences between genotypes within water treatment were analysed using a t-test (*n*=5). Genotypic differences in A/E under specific water treatment were obtained by applying a parallelism test.

### IL20 exhibited higher root influx at an earlier time of day under drought stress

Next, we hypothesized that the ability of IL20 to maintain higher gas exchange under DS, alongside its higher root-to-shoot ratio, may suggest better water extraction from the soil. To test this hypothesis, we analysed the daily and hourly longitudinal root influx of the two genotypes. In general, under both water treatments, IL20 had a higher root influx compared with Svevo, although, under DS, IL20 maintained its root influx rate, as Svevo root influx decreased with intensified water stress (Fig. 5A, Supplementary Fig. S4). To capture the hourly differences during the most intense phase of water stress, we averaged the daily maximum influx per hour for each genotype during the last five days. Under WW, a similar daily pattern was observed between the genotypes where IL20 maintained a higher root influx throughout the day (Fig. 5B). Under DS, IL20 exhibited a higher root influx compared with Svevo, mainly during the morning hours (07:00-12:00) when VPD was relatively lower (Fig. 5C; Supplementary Video S1). This phenomenon can support lower water loss per unit of carbon assimilation under DS, derived from the combination of maximization root influx and gs_c_ in the early morning time when VPD is low.

**Figure 5.**
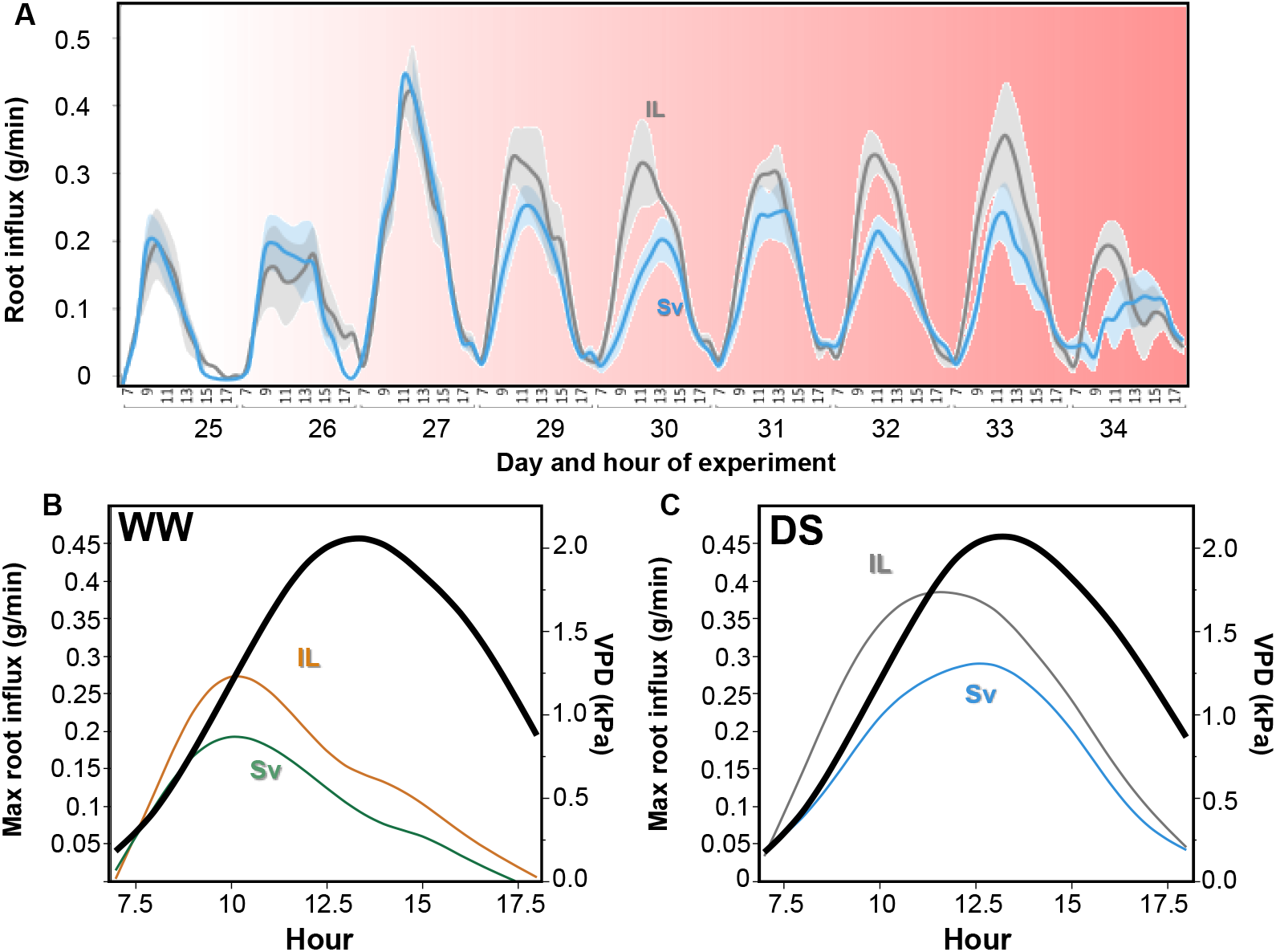
Root influx dynamics of Svevo (Sv) and IL20 (IL) under well-watered (WW) and drought stress (DS) treatments. (**A**) Hourly root influx dynamics throughout the increasing drought intensity. The increasing drought intensity is represented with the pink background colour intensity. The average maximum hourly root influx was calculated for the last 5 days (30-34) of the experiment under (**B**) WW and (**C**) DS treatments. Hourly vapour pressure deficit (VPD) is represented by the black line. The genotype curves represent the means (*n*=5) within water treatment, day and hour. Shaded colours indicate the standard error.

### Structural equation modelling highlights the multidimensional interactions of genotypic-atmospheric-physiological factors under drought stress

We next aimed to integrate the various dependent and independent effects of atmospheric and physiological variables by application of a structural equation modelling (SEM) statistical approach for multiple inter-correlated parameters. We assumed an initial path with two latent variables (atmospheric and physiological), representing a concept that one or more observed variables are presumed to be highly correlated with latent variables. The model fit report indicates that the path we suggested was a good fit (χ=2.80, *P*_Chisq_=0.24, and χ/DF=1.40; Supplementary Table S6) according to the Hu and Bentler (1999) and Lomax and Schumacker (2012) methodology of evaluating models to the suggested path. All nine regressions were found significant, which proves the interrelations of genotype-water availability-atmospheric state (Fig. 6). Physiological and atmospheric endogenous variables had relatively high R^2^ values while the latent variable of physiology had lower values (R^2^=0.23) that may result from the multi-dimensionality of diurnal physiological response to the atmospheric cues (Supplementary Table S7). The genotypic factor (fundamental factor) has a significant effect on physiological traits (latent variable) and VWC. (Supplementary Table S8). Interestingly, VPD and radiation presented opposite effects on plant water intake and loss. While VPD negatively affects transpiration rate, gs_c_ and root influx, radiation had a positive influence. Notably, root influx was less affected by radiation compared with gs_c_ and transpiration rate.

**Figure 6.**
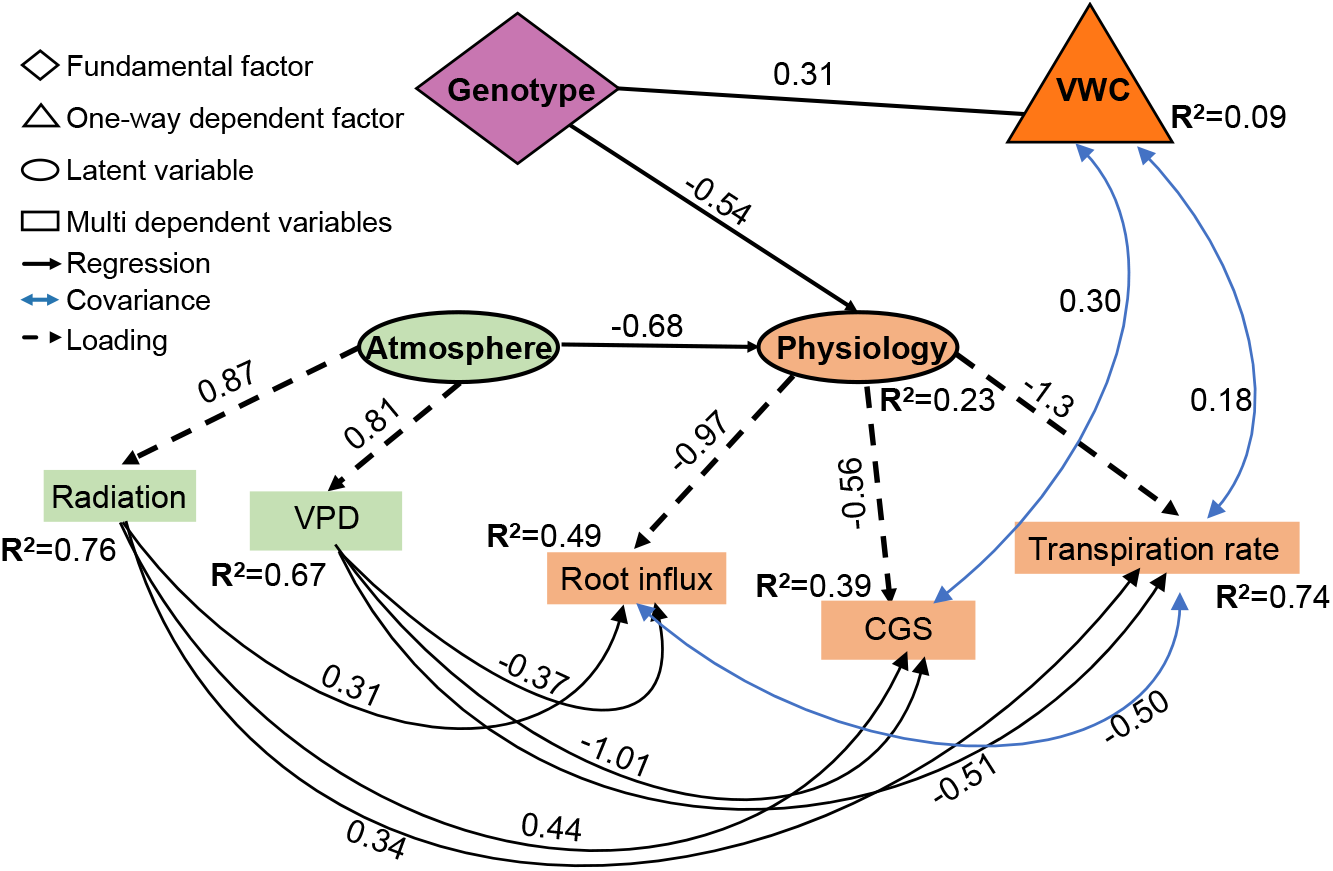
Scheme of the genetic-physiological-atmospheric network path under drought stress. Black and solid one-directional arrows represent standardized parameter estimates for regressions, and dashed arrows represent the loadings standardized parameter estimates. The blue two-directional arrows represent the standardized parameter estimates for covariance and R^2^ for the endogenous variables. Only significant paths are presented (*P*≤0.05).

## Discussion

The unpredictable and fluctuating atmospheric conditions associated with present and projected climate change pose a challenge for wheat cultivation and improvement. Under such conditions, developmental plasticity is an essential adaptive mechanism to promote future production (Gray and Brady, 2016). Phenotypic plasticity in the root-to-shoot ratio can support productiveness under water stress as plant growth potential reduces and root growth is favoured over the shoot to limit evaporation and extract water residuals (Reich, 2002; Correa *et al*., 2019). Wheat breeding has focused on aboveground traits, and root system response to water stress has been less studied. Therefore, new genetic sources for expanding the range of root-to-shoot ratio can be valuable. Recently, we identified a novel mechanism of a drought-induced root-to-shoot shift from wild emmer wheat. Here we harness this adaptive wild introgression line to elucidate the advantage in promoting higher responsiveness to drought and atmospheric state by synchronizing the diurnal water usage. This soil–plant–atmosphere continuum response results in higher transpiration use efficiency and mediates the wheat water influx continuum under drought stress.

Our findings show that the elite cultivar Svevo exhibited a significant reduction in shoot and root DW [−12% (*P*=0.055) and −41% (*P*=0.0003), respectively] which resulted in a 33% reduction of root-to-shoot ratio. In contrast, IL20 showed a milder reduction in shoot DW (−8%; *P*=0.18) and non-significant increment in root DW (+11%, *P*=0.11), which resulted in a 21% increase in root-to-shoot ratio (Fig. 1). Accordingly, IL20 exhibited significantly greater whole-canopy transpiration and root influx (Figs. 2 and 5). It has been suggested that under rainfed Mediterranean agro-systems, limiting the vegetative growth (*i.e*., transpiration rate) is a desirable trait as it can preserve more soil moisture for the reproductive phase (Passioura, 1983). Therefore, we can expect that the higher biomass of IL20 will have a negative effect during the progression of stress and increasing drought intensity. However, while Svevo showed a significant reduction in biomass accumulation with increasing stress intensity, IL20 was able to maintain growth (Fig. 1A-B). This advantage of IL20 is expressed in its ability to maintain greater transpiration and gs_c_ under DS (Figs. 2, 3). The plant’s ability to maintain higher transpiration under DS can distinguish between tolerant and susceptible wheat cultivars (El Habti *et al*., 2020). These results, as well as our findings, highlight the important role of maintaining transpiration under drought stress to support wheat productiveness.

Under semi-arid conditions, transpiration efficiency (TE) is a key adaptive trait (Collins *et al*., 2021), that can be mediated by stomatal responsiveness to environmental cues. For example, maximizing the transpiration rate under a preferable atmospheric state has been suggested as a breeding target for drought-prone maize (*Zea mays*) agro-system (Messina *et al*., 2015). Likewise, assimilation and transpiration rates declined when atmospheric VPD increased in drought-adapted soybean (*Glycine max*) genotypes (Fletcher *et al*., 2007). Moreover, a genetic variation between elite wheat cultivars in their transpiration response to given evaporative demand has been reported (Schoppach and Sadok, 2012), which points to the genetic control of TE. Under WW treatment both genotypes exhibited similar gs_c_ before VPD reached its peak (*i.e*., mid-day), whereas, under DS, IL20 exhibited greater gs_c_ early in the morning (07:00-10:00 AM) compared with Svevo (Fig. 3C). This advantage of IL20 under lower VPD promotes higher TE. In addition, IL20 increased its WUE_*l*_ in response to DS. These results are in agreement with Leakey *et al*. (2019), suggesting that a higher response of WUE_*l*_ to VPD may indicate diurnal/spatial sensitivity to environmental factors, although the physiological source of this hyper-responsiveness (rather in the shoot or root tissue) is not yet known. The ability of plants to support higher gas exchange under low VPD during a preferable time window (*i.e*., early morning), could be coupled with increasing water potential and could promote root growth under drought (Quarrie and Jones, 1979). IL20’s higher maximal root influx under both water treatments (Fig. 5A, Supplementary Fig. S3), as expressed also in its advantage in transpiration over Svevo (Fig. 2A), is associated with its larger root and shoot biomass, but does not explain the diurnal root water uptake differences between the genotypes. Characterizing these trends suggests that while IL20 was able to maintain its maximal root influx early in the morning, Svevo’s root influx was delayed and reached its peak an hour later coinciding with the atmospheric VPD peak (Fig. 5B, C). This early water flux could enable IL20 to adjust its diurnal transpiration pattern to the atmospheric conditions, which would result in higher TE under DS.

Schoppach *et al*. (2013) found that drought-tolerant wheat breeding line (cv. RAC875) limited its transpiration rate in response to increasing VPD via smaller xylem vessel anatomy that enables water conservation. Thus suggesting a connection between root anatomy and transpiration regulation. In contrast, a comparison of the wild introgressions with previous genomic dissection of wild emmer QTLs conferring root axial conductance and xylem anatomy (Hendel *et al*., 2021) did not show any overlap. Thus, it suggests that the genetic regulation adapting the gas exchange to favourable climatic periods is associated with root morphology, rather than anatomical modifications. Overall, this wild mechanism offers a new strategy to improve TE by incorporating morphological root-to-shoot ratio increment and VPD sensing mechanism under drought stress. Moreover, we suggest that this distinctive combination is not associated with the well-accepted penalty of increasing TE by reducing the plant size under drought (see Vadez *et al*., 2014).

Previous studies have mainly focused on aboveground tissues, while belowground responsiveness to atmospheric cues has received less attention. To better understand the intra- and inter-relations between plant tissues and atmospheric parameters, we applied the structural equation modelling (SEM) approach. Segmentation of the physiological responses to drought stress indicates that leaf tissue is more responsive to the atmospheric state as expressed in gas exchange alterations. In contrast, the root influx was mostly affected by the VPD, presumably in response to the shoot stomatal closure since VPD had the most major negative effect on gs_c_ (Standardized estimate = −1.01), which resulted in lowering the transpiration rate and finally decreasing root influx. These results are somewhat opposite the daily root hydraulics oscillations and photosynthesis to promote water stress tolerance (Caldeira *et al*., 2014; Tardieu *et al*., 2015). Our SEM analysis, pointing to the leaf as the tissue that responds to the daily atmospheric pattern, affects the root water influx and alters TE. Segmentation of the environmental cue associated with the physiological response to water stress is still puzzling since the co-occurrence of VPD and radiation makes it challenging to untangle their effects (Grossiord *et al*., 2020; Fig. 3). Our model enabled us to dissect these two parameters although they showed similar daily patterns (Fig. 3). VPD was the major parameter that controlled plant water flux, with a negative effect on the physiological performance, while radiation had a positive effect that increases gs_c_, transpiration rate and root influx (Supplementary Table S8). These robust data aggregations enable modelling the inter-relations between atmospheric cues and physiological response under increasing water shortage.

### Concluding remarks

Under the semi-arid Mediterranean basin conditions, breeding drought-tolerant cultivars is based on their high sensitivity to atmospheric changes (Sadok and Tamang, 2019). Lopez *et al*., (2020) suggested that the most useful approach for understanding the VPD effect on plant physiological response will simultaneously examine traits that are expressed in the different organs and will link the physiological processes on a larger scale, based on models. We integrated these approaches and suggested that plants’ adaptation to water and atmospheric stress is a combination of physiological response (alteration in root-to-shoot ratio) and increased transpiration efficiency by matching the plant water in/out fluxes to the early morning. Modelling the root and shoot tissue response to the atmospheric state indicated that breeding drought-tolerant (water and atmospheric) cultivars should be focused on leaf hypersensitive response to VPD and a more efficient root system that can increase influx and support higher gas exchange during suitable atmospheric conditions.

## Abbreviations

A: Assimilation rate
DS: drought stress
DW: dry weight
IL: introgression line
E: normalized transpiration rate
gs: stomatal conductance
SEM: structural equation modelling
VPD: vapour pressure deficit
WUE: water-use efficiency
WW: well-watered
gs_c_: whole canopy conductance

## Supplementary material

**Fig. S1.** Daily and hourly atmospheric data throughout the experiment.

**Fig. S2.** Flag leaf stomatal number under well-watered and drought stress treatments.

**Fig. S3.** Leaf total conductance to water vapour (g_tw_) of Svevo and IL20 under well-watered (WW) and drought stress (DS) treatments.

**Fig. S4.** Hourly root influx dynamics for the last nine days of the experiment under well-watered treatment.

**Video S1** Daily diurnal root influx dynamics of Svevo and IL20 under drought stress treatment.

**Table S1.** The longitudinal growth rate of Svevo and IL20 under well-watered and drought stress treatments.

**Table S2.** Morpho-physiological traits on the last day of the experiment.

**Table S3.** Longitudinal daily transpiration of Svevo and IL20 under well-watered and drought stress treatments.

**Table S4.** Leaf gas exchange and water use efficiency under well-watered and drought stress treatments.

**Table S5.** Leaf water-use efficiency fit model and parallelism test.

**Table S6.** Structural equation model fit report.

**Table S7.** R^2^ for the structural equation model endogenous variables.

**Table S8.** Structural equation model descriptive statistics. Standardized estimate variables under drought stress.

## Acknowledgments

We thank members of the Peleg lab for their helpful comments during the preparation of this manuscript. We thank Dr. R. Hayouka and E. Hendel for technical assistance with the experiments and Mrs. L. Shemesh for illustrating the wheat phenological stages. This study was partially supported by the State of Israel Ministry of Agriculture and Rural Development (grants # 20-10-0066; 12-01-0005), the U.S. Agency for International Development Middle East Research and Cooperation (grant # M34-037), and the Dutch Ministry of Foreign Affairs under Dutch development / foreign policy (Project WheatMAX).

