## Supplementary material for "Modifying root/shoot ratios improves root water influxes in wheat under drought stress": SI Figures

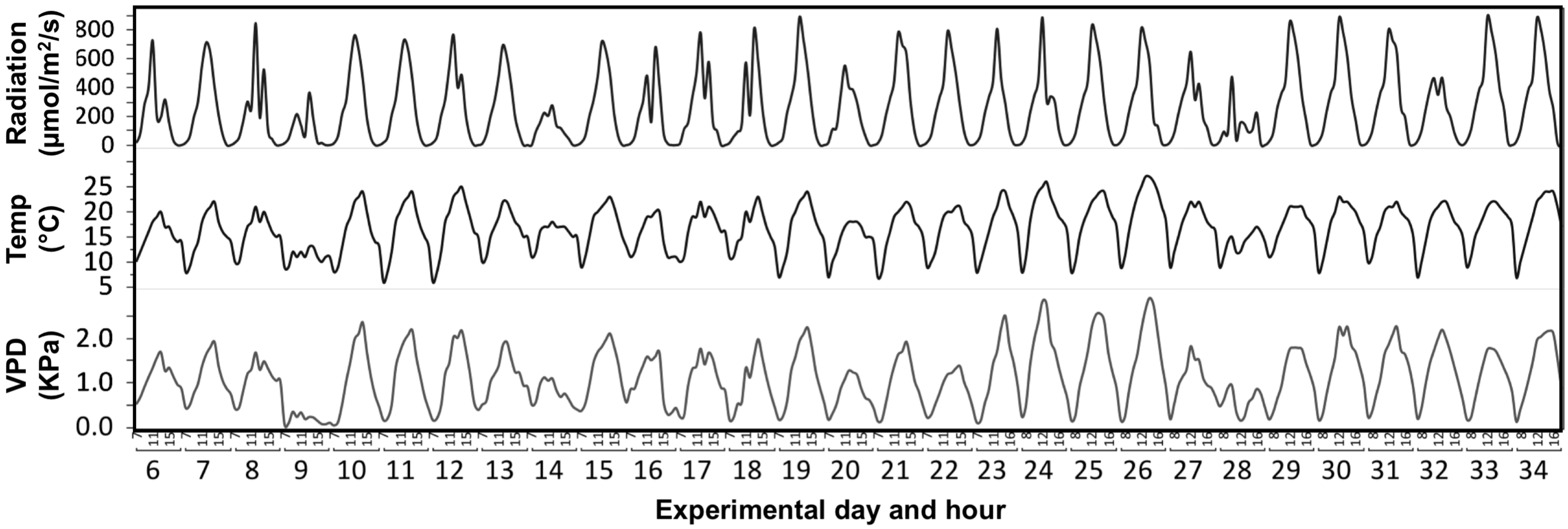


**Supplementary Fig. 1.** Daily and hourly atmospheric data along the experiment. Photosynthetically active radiation (Radiation), temperature (Temp) and vapour pressure deficit (VPD).

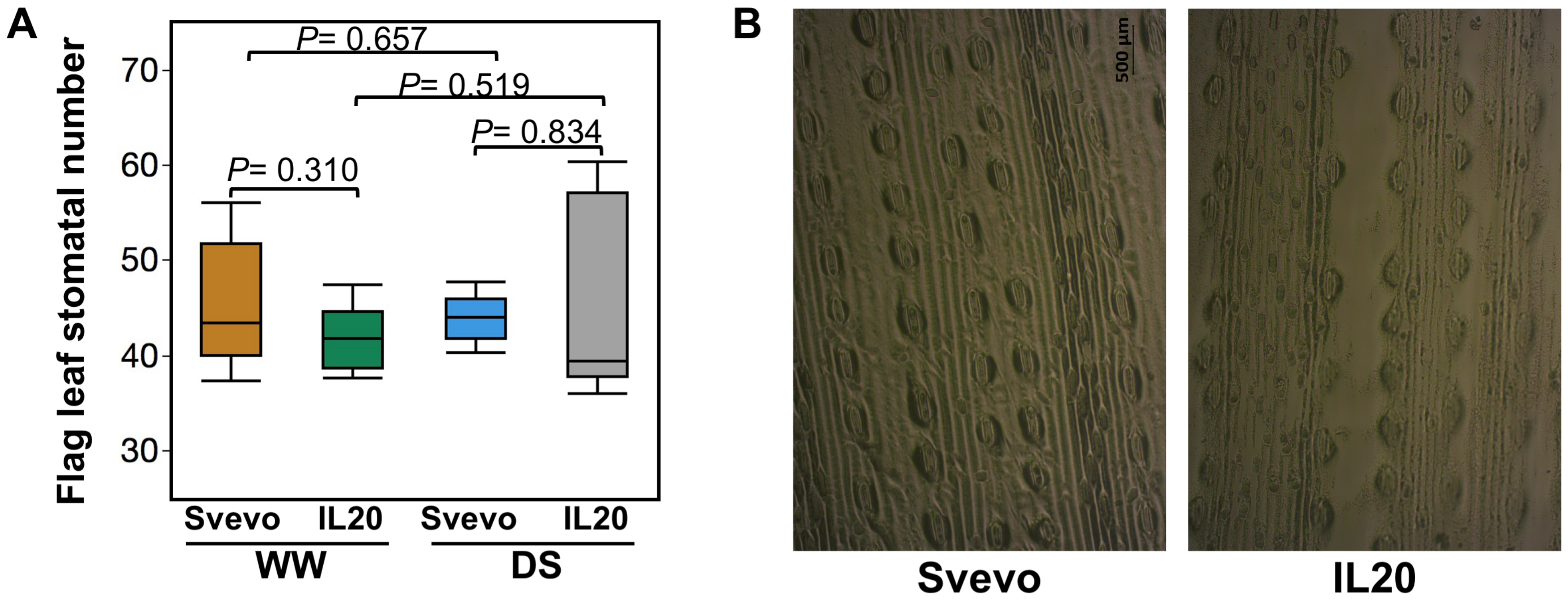


**Supplementary Fig. S2.** Flag leaf stomata number of Svevo and IL20 under contrasting water treatments. (**A**) Flag leaf stomatal number under well-watered (WW) and drought stress (DS) treatments in 0.2 $cm^2$ leaf. (**B**) A representative image of stomatal pattern in adaxial side of the flag leaf. Differences between genotypes within water treatment were analysed using a t-test (*n*=4).

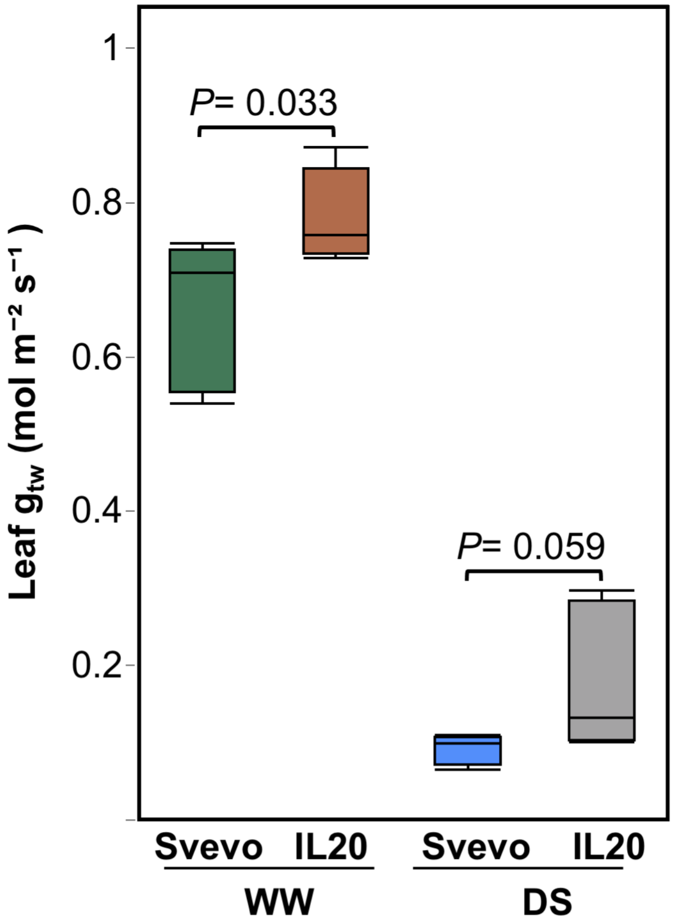


**Supplementary Fig. S3.** Leaf total conductance to water vapour ($g_{tw}$) of Svevo and IL20 under well-watered (WW) and drought stress (DS) treatments. Differences between genotypes within water treatment were analysed using a t-test (*n*=5).

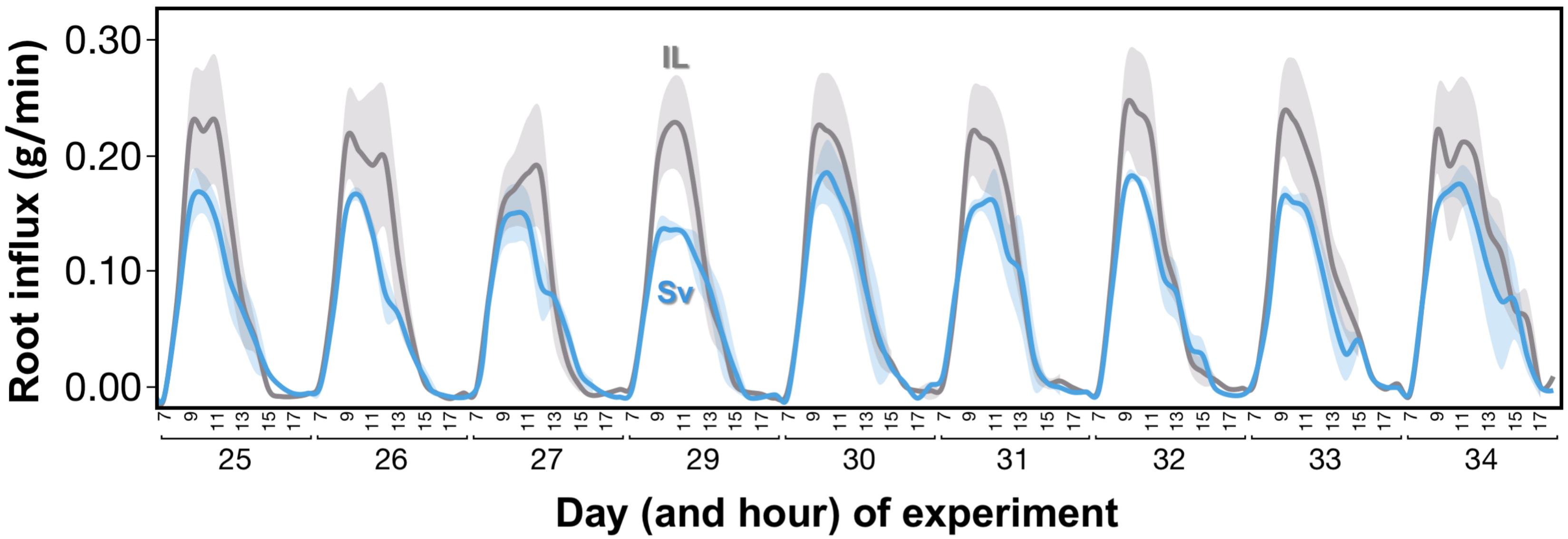


**Supplementary Fig. S4.** Hourly root influx dynamics of Svevo (Sv, blue) and IL20 (IL, grey) at the last nine days of the experiment under well-watered treatment (*n*=5) .
